## Supplementary material for "Integrative QTL analysis of gene expression and chromatin accessibility identifies multi-tissue patterns of genetic regulation": File S1

### Data and Supplement details

Files in the **Supplement** include:

- File_S1: `data_supplement_details.pdf`
- Results files:
  - File_S2 - S4: `de_{liver_lung,liver_kidney,lung_kidney}.csv`
  - File_S5 - S7: `dar_{liver_lung,liver_kidney,lung_kidney}.csv`
  - File_S8 - S10: `{liver,lung,kidney}_local_eqtl.csv`
  - File_S11 - S13: `{liver,lung,kidney}_distal_eqtl.csv`
  - File_S14 - S16: `{liver,lung,kidney}_local_cqtl.csv`
  - File_S17 - S19: `{liver,lung,kidney}_distal_cqtl.csv`
  - File_S20 - S22: `{liver,lung,kidney}_chromatin_mediation.csv`
- Additional resource files:
  - File_S23: `supplement_tables_figures.pdf`
  - File_S24: `appendices.pdf`
- Software files:
  - File_S25: `miqtl_1.1.2.tar.gz`
- R code:
  - File_S26: `generate_manuscript_figures.R`
  - File_S27: `generate_supplement_tables_figures.R`
  - File_S28: `plotting_functions.R`
  - File_S29: `analysis_functions.R`
  - File_S30: `make_qtl_tables_functions.R`
  - File_S31: `miqtl_code_demonstration.R`

The data necessary for these analyses and raw QTL results files have been stored at figshare (doi:10.6084/m9.figshare.9985514). These files include:

- Data files:
  - `cc_genome_cache_full_l2_0.1.zip`
    - Founder haplotype mosaics for the 47 strains based on Build37, derived from files at `http://csbio.unc.edu/CCstatus/index.py?run=FounderProbs`, used for haplotype-based QTL analysis. Haplotype intervals were thinned, averaging adjacent intervals for which the mean l2 norm < 0.1 [1]. Data are arranged into a directory tree structure (genome cache) that conforms to the HAPPY format [2].
  - `{liver,lung,kidney}_expression.csv.zip`
  - `{liver,lung,kidney}_chromatin.csv.zip`
  - `refseq_mm9_tss.txt.zip`
    - Gene annotation data for Build37, providing gene transcription start site information necessary for determining local/distal status for eQTL.
  - `pik3c2g_phylogeny.csv`

    * Maps the founder haplotypes around *Pik3c2g* to the three *Mus* sub-species lineages (`http://msub.csbio.unc.edu`) [3].
  - `isvdb_var_db.tar.gz`
    * SNP dosages derived from the founder haplotype mosaics and variant-to-strain distribution pattern, using ISVdb (`http://isvdb.unc.edu`) [4].
  - `qtl_intervals_for_var_association.csv`
    * Defined regions around detected QTL used to select variants for association analysis.
- Raw results files:
  - `raw_{liver,lung,kidney}.eqtl_methodG.csv`
  - `raw_{liver,lung,kidney}.eqtl_methodL.csv`
  - `raw_{liver,lung,kidney}.eqtl_methodC.csv`
  - `raw_{liver,lung,kidney}.cqtl_methodG.csv`
  - `raw_{liver,lung,kidney}.cqtl_methodL.csv`
  - `raw_{liver,lung,kidney}.cqtl_methodC.csv`

### miQTL package

The static version of the miQTL R package (1.1.1) used for this work is provided in the **Supplement** as File_S2. The current version of miQTL is available here: `https://github.com/gkeele/miqtl`. miQTL can be installed using the command 'R CMD INSTALL' at the terminal. The current version from GitHub can be conveniently installed using the devtools R package and the following command within R: 'install_github("gkeele/miqtl")'.

### File types

- *.csv - Comma-separated value files representing the trait data used for differential and QTL analyses and their large results tables, which were too large to be included as formatted tables.
- *.R - R scripts used to run the statistical analyses and generate the figures.
- *.RData - The zipped directory contained in `cc_genome_cache_full_l2_0.1.zip` is composed of *.RData files that are HAPPY formatted, which the miQTL R package is designed to use.
- *.txt - The zipped text file `refseq_mm9_tss.txt.zip` is tab-delimited.
