## Supplementary material for "Integrative QTL analysis of gene expression and chromatin accessibility identifies multi-tissue patterns of genetic regulation": File S23

### Supplemental tables and figures for Keele & Quach *et al.*

Table S1: **Number of differentially expressed genes and accessible chromatin regions detected between liver, lung, and kidney tissues at FDR** $\leq 0.1$

| | | Tissue comparison | | |
|---|---|---|---|---|
| | | Liver/lung | Liver/kidney | Lung/kidney |
| Genes | Up-regulated | 2,473 (20.8[a]) | 2,123 (17.9[a]) | 2,246 (18.9[a]) |
| | Down-regulated | 3,236 (27.2[a]) | 1,441 (12.1[a]) | 2,527 (21.3[a]) |
| | Total | 5,709 (48.0[a]) | 3,564 (30.0[a]) | 4,773 (40.2[a]) |
| Chromatin regions | Increased accessibility | 20,194 (19.6[b]) | 15,252 (12.9[b]) | 19,202 (17.4[b]) |
| | Decreased accessibility | 20,603 (19.7[b]) | 12,796 (11.4[b]) | 12,967 (11.2[b]) |
| | Total | 40,797 (39.3[b]) | 28,048 (24.3[b]) | 32,169 (28.6[b]) |

[a] Percentage of all tested genes.
[b] Percentage of all tested chromatin regions prior to merging adjacent genomic windows.

Table S2: **Number of genes with eQTL detected in liver, lung, and kidney tissues at FDR** $\leq 0.1$

| | | Tissue (%) | | |
|---|---|---|---|---|
| Procedure | eQTL type | Liver | Lung | Kidney |
| Analysis G | All | 520 (6.2[a]) | 478 (4.2[a]) | 739 (7.3[a]) |
| | Local[d] | 400 (76.9[b]) | 369 (77.2[b]) | 601 (81.3[b]) |
| | Distal[e] | 132 (25.4[b]) | 112 (23.4[b]) | 148 (20.0[b]) |
| Analysis C | All | 2,587 (30.8[a]) | 2,069 (18.2[a]) | 3,191 (31.6[a]) |
| | Local[d] | 1,749 (67.6[c]) | 1,498 (72.4[c]) | 2,214 (69.4[c]) |
| | Distal[e] | 838 (32.4[c]) | 571 (27.6[c]) | 977 (30.6[c]) |
| Analysis L[f] | Genome-wide FWER $< 0.05$ | 702 (8.4[a]) | 713 (6.3[a]) | 955 (9.5[a]) |
| | Chromosome-wide FWER $< 0.05$ | 1,661 (19.8[a]) | 1,880 (16.6[a]) | 2,102 (20.8[a]) |

[a] Percentage of all tested genes.
[b] Percentage of genes with eQTL from Analysis G.
[c] Percentage of genes with eQTL from Analysis C.
[d] Within 10Mb upstream or downstream of gene TSS.
[e] More than 10Mb upstream or downstream of gene TSS, or on another chromosome.
[f] Not FDR controlled.

Table S3: **Number of genes with eQTL detected in liver, lung, and kidney tissues at FDR** $\leq 0.2$

| Procedure | eQTL type | Tissue (%) Liver | Lung | Kidney |
|---|---|---|---|---|
| Analysis G | All | 881 (10.5[a]) | 811 (7.1[a]) | 1,189 (11.8[a]) |
| | Local[d] | 568 (64.5[b]) | 522 (64.4[b]) | 809 (68.0[b]) |
| | Distal[e] | 339 (38.5[b]) | 301 (37.1[b]) | 411 (34.6[b]) |
| Analysis C | All | 4,699 (55.9[a]) | 3,675 (32.4[a]) | 5,469 (54.2[a]) |
| | Local[d] | 2,519 (53.6[c]) | 2,213 (60.2[c]) | 3,099 (56.7[c]) |
| | Distal[e] | 2,180 (46.4[c]) | 1,462 (39.8[c]) | 2,370 (43.3[c]) |

[a] Percentage of all tested genes.
[b] Percentage of genes with eQTL from Analysis G.
[c] Percentage of genes with eQTL from Analysis C.
[d] Within 10Mb upstream or downstream of gene TSS.
[e] More than 10Mb upstream or downstream of gene TSS, or on another chromosome.

Table S4: **Number of chromatin accessibility sites with cQTL detected in liver, lung, and kidney tissues at FDR** $\leq 0.1$

| Procedure | cQTL type | Tissue (%) Liver | Lung | Kidney |
|---|---|---|---|---|
| Analysis G | All | 17 (0.1[a]) | 114 (0.5[a]) | 59 (0.3[a]) |
| | Local[d] | 16 (94.1[b]) | 78 (68.4[b]) | 39 (66.1[b]) |
| | Distal[e] | 1 (5.9[b]) | 36 (31.6[b]) | 20 (33.9[b]) |
| Analysis C | All | 39 (0.3[a]) | 226 (0.9[a]) | 130 (0.7[a]) |
| | Local[d] | 35 (89.7[c]) | 186 (82.3[c]) | 98 (75.4[c]) |
| | Distal[e] | 4 (10.3[c]) | 40 (17.7[c]) | 32 (24.6[c]) |
| Analysis L[f] | Genome-wide FWER $< 0.05$ | 70 (0.6[a]) | 244 (1.0[a]) | 149 (0.8[a]) |
| | Chromosome-wide FWER $< 0.05$ | 299 (2.6[a]) | 876 (3.6[a]) | 616 (3.4[a]) |

[a] Percentage of all tested chromatin regions.
[b] Percentage of genes with cQTL from Analysis G.
[c] Percentage of genes with cQTL from Analysis C.
[d] Within 10Mb upstream or downstream of chromatin region midpoint.
[e] More than 10Mb upstream or downstream of chromatin region midpoint, or on another chromosome.
[f] Not FDR controlled.

Table S5: **Number of chromatin accessibility sites with cQTL detected in liver, lung, and kidney tissues at FDR** $\leq 0.2$

| | | Tissue (%) | | |
|---|---|---|---|---|
| Procedure | cQTL type | Liver | Lung | Kidney |
| Analysis G | All | 20 (0.2[a]) | 220 (0.9[a]) | 113 (0.6[a]) |
| | Local[d] | 18 (90.0[b]) | 116 (52.7[b]) | 55 (48.7[b]) |
| | Distal[e] | 2 (10.0[b]) | 105 (47.7[b]) | 58 (51.3[b]) |
| Analysis C | All | 62 (0.5[a]) | 309 (1.3[a]) | 249 (1.4[a]) |
| | Local[d] | 50 (80.6[c]) | 238 (77.0[c]) | 149 (59.8[c]) |
| | Distal[e] | 12 (19.4[c]) | 71 (23.0[c]) | 100 (40.2[c]) |

[a] Percentage of all tested chromatin regions.
[b] Percentage of genes with cQTL from Analysis G.
[c] Percentage of genes with cQTL from Analysis C.
[d] Within 10Mb upstream or downstream of chromatin region midpoint.
[e] More than 10Mb upstream or downstream of chromatin region midpoint, or on another chromosome.

Table S6: **Number of genes with chromatin mediation of local-eQTL in liver, lung, and kidney tissues**

| | | Tissue (%) | | |
|---|---|---|---|---|
| Procedure | Significance | Liver | Lung | Kidney |
| Analysis L[b] | genome-wide | 13 (0.8[a]) | 42 (2.2[a]) | 21 (1.0[a]) |
| | chromosome-wide | 35 (2.1[a]) | 106 (5.6[a]) | 66 (3.1[a]) |

[a] Percentage of genes with a chromatin mediator that meet chromosome-wide significance for a local-eQTL through Analysis L.
[b] 10Mb upstream or downstream of the gene TSS with local-eQTL that meets chromosome-wide significance through Analysis L. Additionally, chromatin mediator must possess a local-cQTL that meets chromosome-wide significance through Analysis L.

Table S7: **Genes with distal-eQTL with gene mediators detected in lung and kidney tissues**

| Tissue | Gene | Chr | Mediator gene | Chr | permP$_{\mathrm{G}}^{m}$ |
|---|---|---|---|---|---|
| Lung | *Akr1e1* | 13 | *Zfp985* | 4 | 4.70e-07 |
| | *Ccnyl1* | 1 | *Zfp979* | 4 | 1.71e-10 |
| | | | *Zfp985* | 4 | 1.23e-05 |
| | *Man2c1* | 9 | *Vash1* | 12 | 1.77e-06 |
| | *Rbm46* | 3 | *Gatm* | 2 | 1.06e-02 |
| Kidney | *C330018D20Rik* | 13 | *Depdc1b* | 18 | 9.97e-04 |
| | *Fbln5* | 12 | *Crym* | 7 | 3.67e-02 |
| | *Gcdh* | 8 | *Dmgdh* | 13 | 8.96e-04 |
| | *Oscp1* | 4 | *Slc25a34* | 4 | 3.27e-03 |

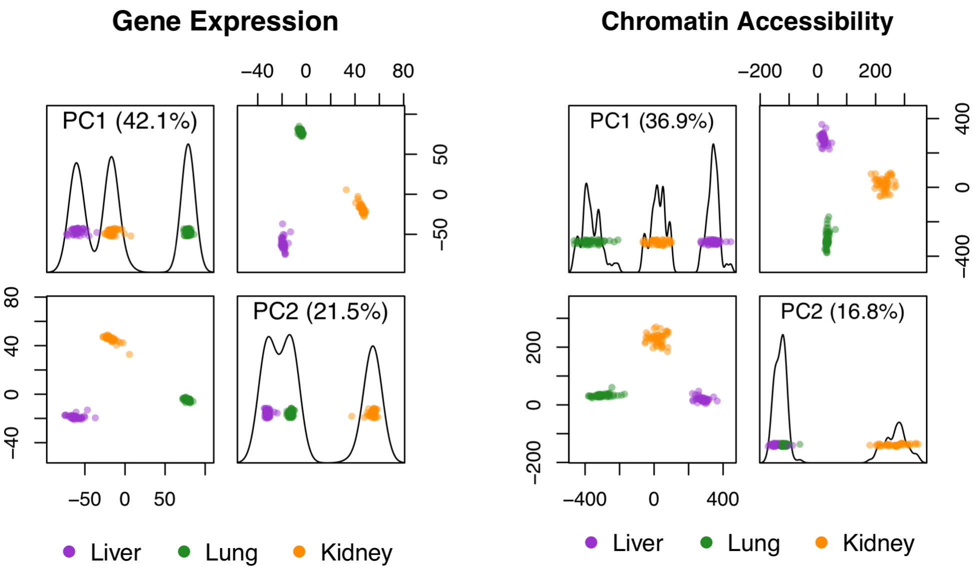


Fig S1: **Principle components analysis identifies tissue type as key source of variation for gene expression and chromatin accessibility.** Molecular traits for liver (purple), lung (green), and kidney (orange) tissue samples were derived from RNA-seq and ATAC-seq data. Principal components (PC) 1 and 2 capture a majority of the variation and show a greater amount of between tissue variability than within tissue variability.

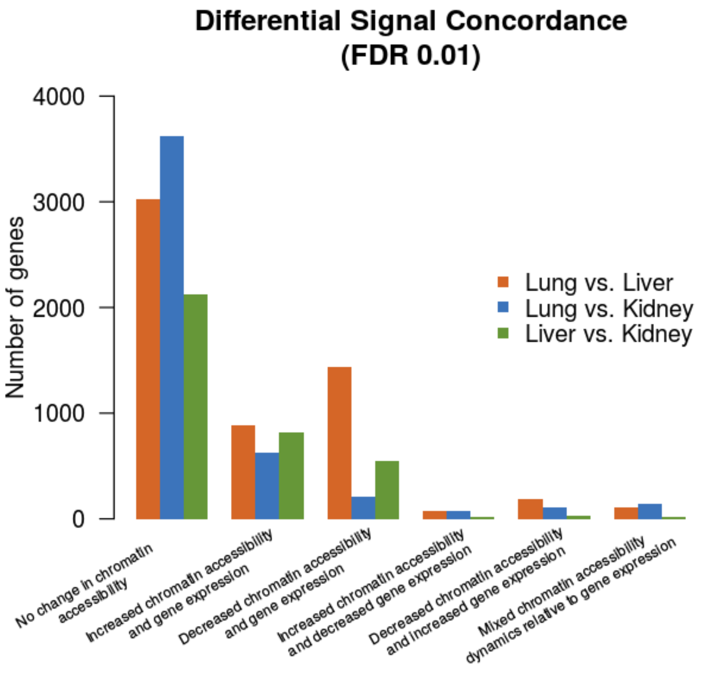


Fig S2: **Concordance between differentially expressed genes and differentially accessible regions in between-tissue comparisons.** Genes were categorized by the direction of the difference in expression and chromatin accessibility in their promoter regions.

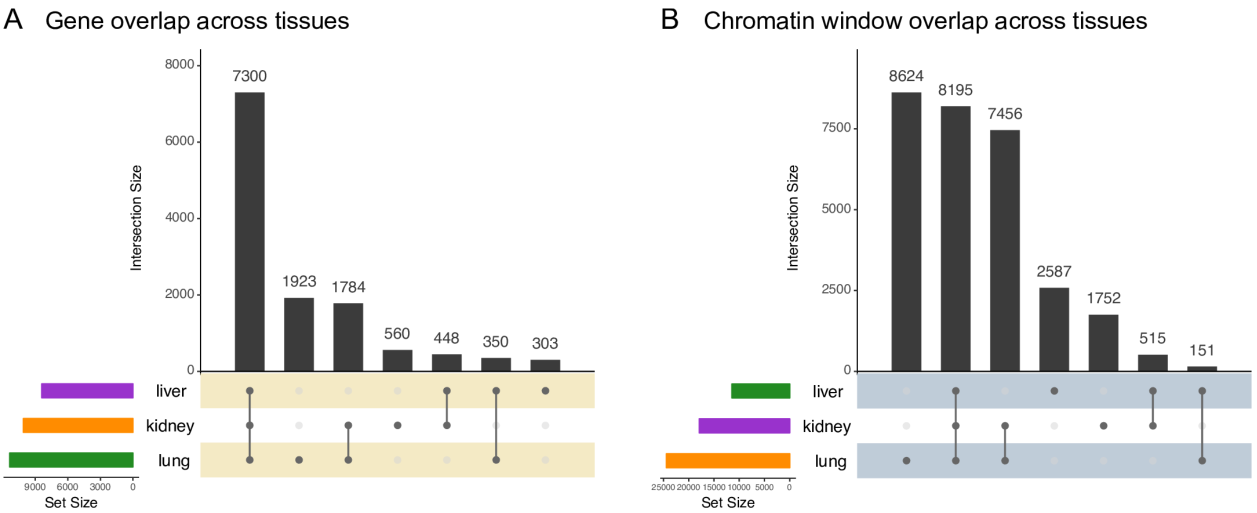


Fig S3: **Overlap across tissues of (A) genes and (B) chromatin windows used for QTL analysis.** Sequence traits were filtered to remove outcomes more likely to cause spurious QTL signals. Genes with TPM $\leq 1$ and chromatin windows with TMP $\leq 5$ for $\geq$ 50% of samples were removed from analysis. After this filtering process, lung had the greatest number of traits analyzed, for both genes and chromatin windows, followed by kidney and then liver.

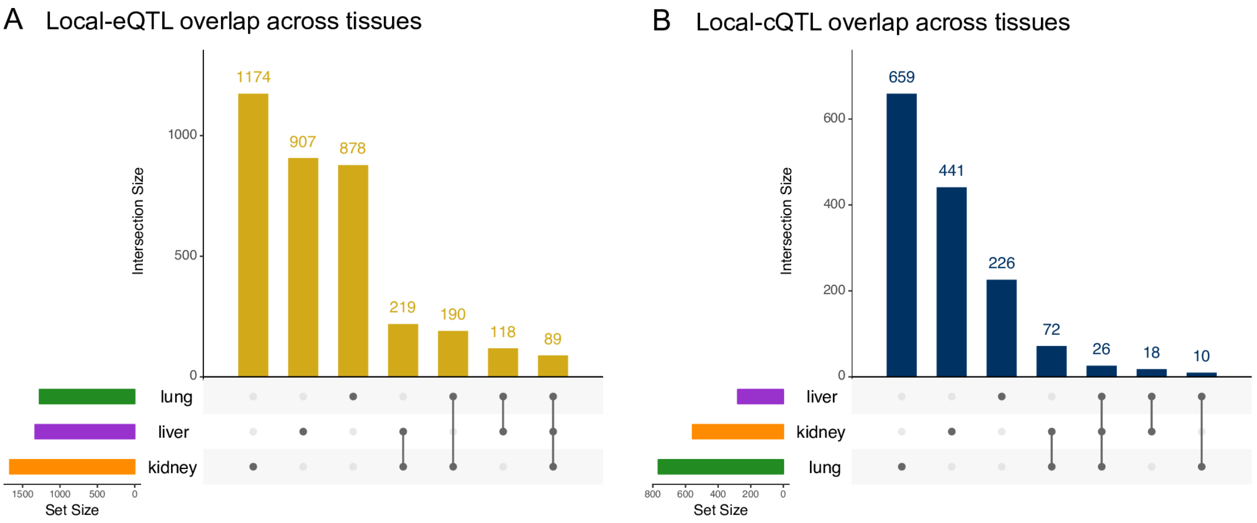


Fig S4: **Overlap across tissues of (A) genes and (B) chromatin windows with local-QTL detected.** The majority of sequence traits with a local-QTL detected were identified in only a single tissue. Kidney had the highest number of local-eQTL, whereas lung had the highest number of local-cQTL. Liver had a relative lack of local-cQTL, which may relate to its having the fewest chromatin windows analyzed (Fig S3B). Results included local-QTL detected with Analysis G (FDR $\leq 0.1$), Analysis C (FDR $\leq 0.1$), and Analysis L (genome-wide and chromosome-wide).

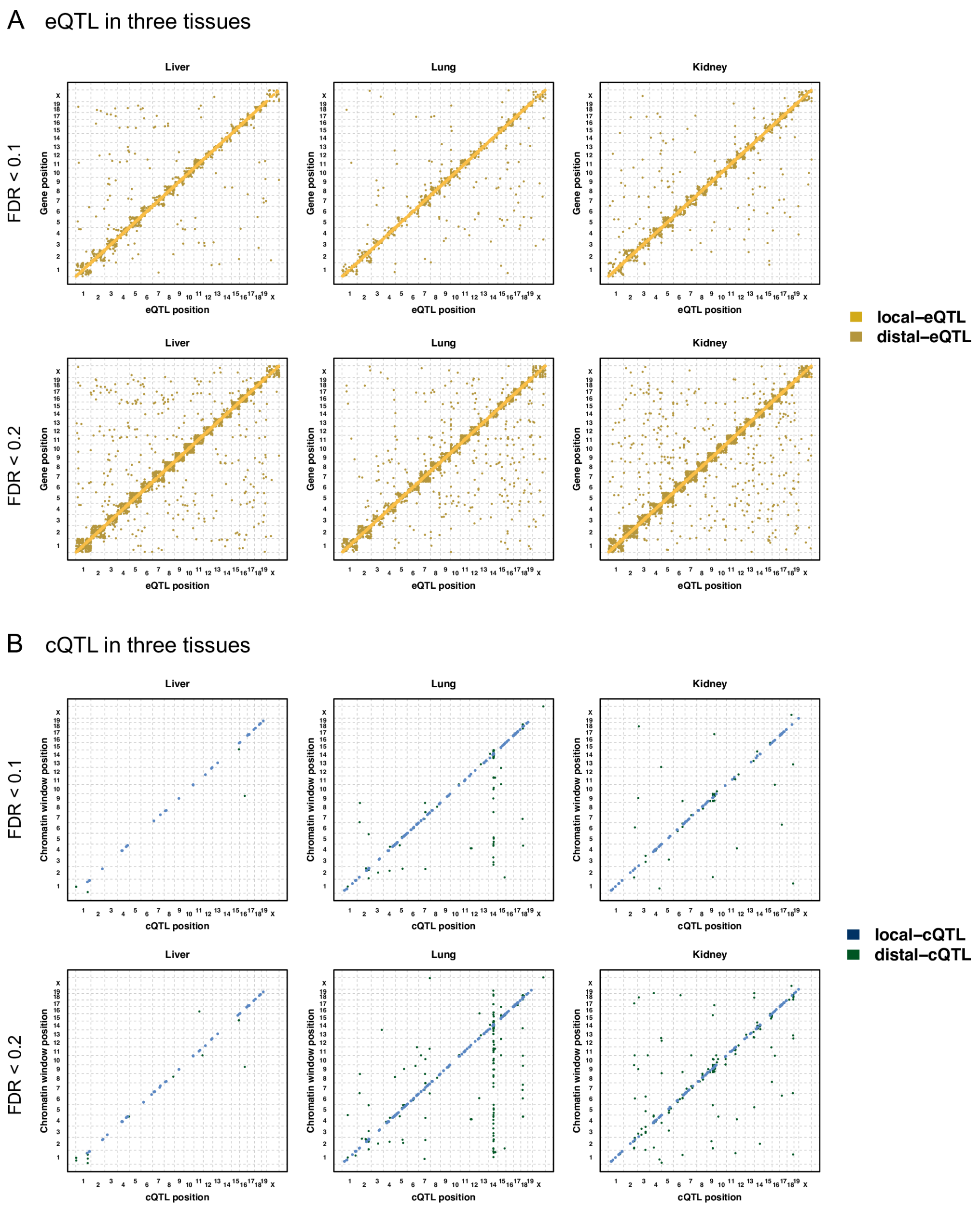


Fig S5: **QTL mapping results using only Analysis G or Analysis C.** QTL map plots of (A) eQTL and (B) cQTL with FDR controlled at 0.1 and 0.2 for liver, lung, and kidney. Detected QTL from Analysis G (multi-stage FDR) and Analysis C (chromosome-wide FDR) are included. Analysis C, which uses FDR control for chromosome-wide significant QTL, produces a large number of intra-chromosomal distal-QTL. The y-axis represents the genomic position of the gene or chromatin site, and the x-axis represents the genomic position of the QTL. Local-QTL appear as dots along the diagonal.

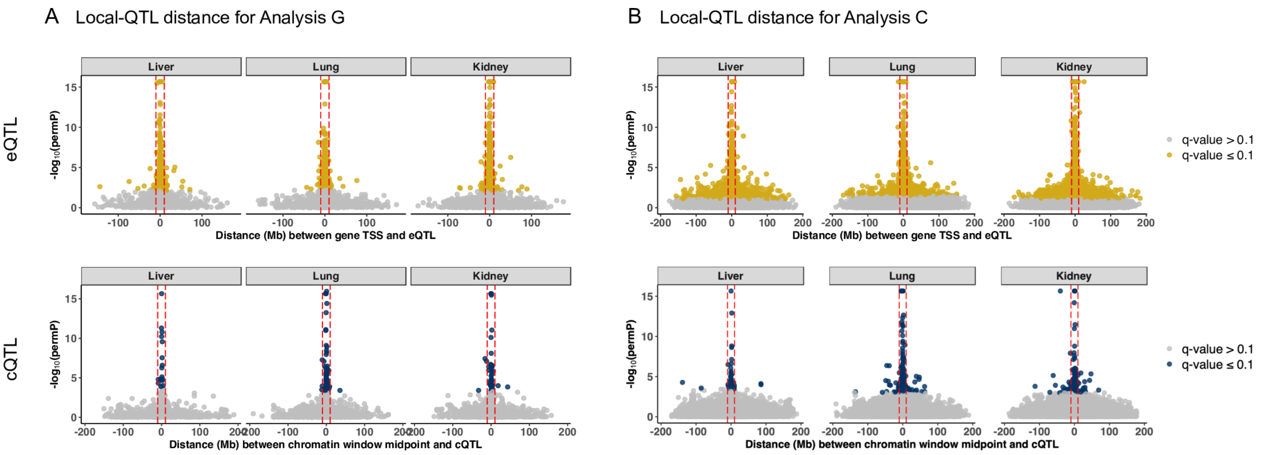


Fig S6: **Highly significant QTL map nearby the gene TSS and chromatin window midpoint.** The permutation-based $p$-value (permP) from (A) Analysis G and (B) Analysis C for eQTL and cQTL by their distance (Mb) from the gene TSS and the midpoint of the chromatin site. Inter-chromosomal distal-QTL are not included. The red dashed lines represent $\pm$10Mb of the gene TSS or the midpoint of the chromatin site for classifying QTL as local or distal. Significant signals (yellow or blue), based on FDR $\leq 0.1$, are largely local. Analysis C detects many more intra-chromosomal distal-QTL.

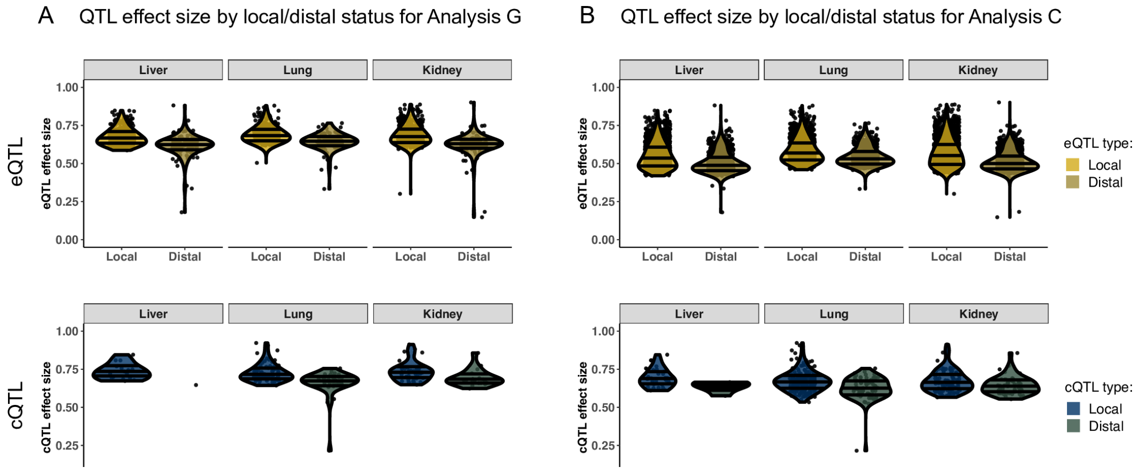


Fig S7: **QTL effect size by local/distal status.** Each dot represents a QTL detected through either (A) Analysis G or (B) Analysis C with FDR $\leq 0.1$. The three horizontal bars represent the 25$^{th}$, 50$^{th}$, and 75$^{th}$ quantiles of QTL effect sizes for all local-QTL per tissue. More local-eQTL are detected and have higher effects than distal-QTL. Analysis C detects a large number of intra-chromosomal distal-QTL that Analysis G does not, many of which have low effect sizes. Effect size estimates are based on a fixed effects model.

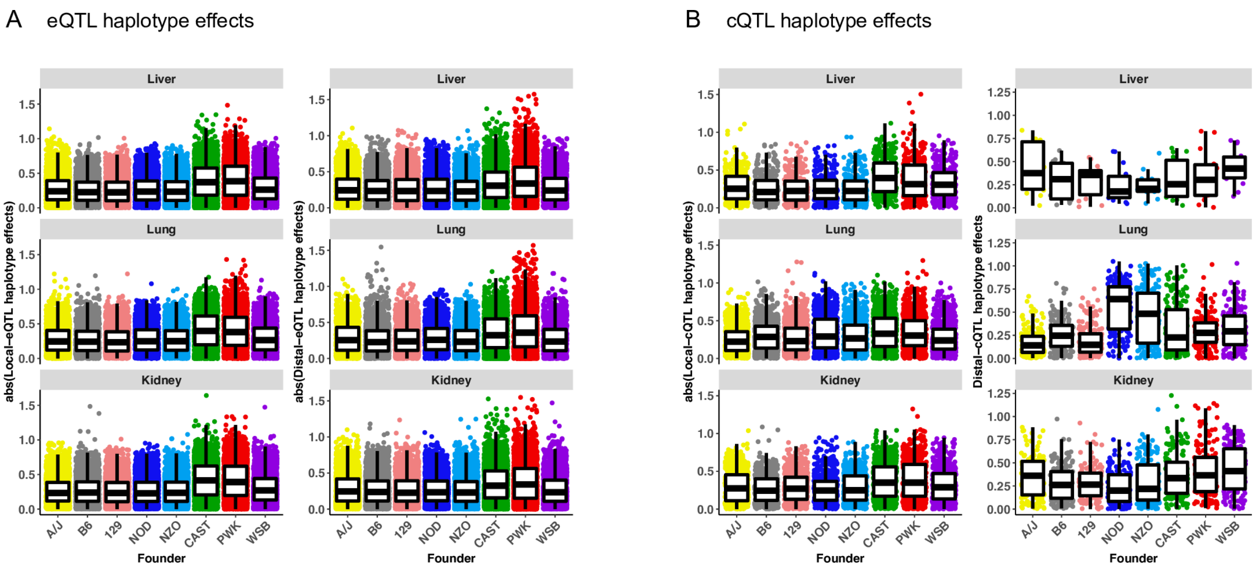


Fig S8: **CAST and PWK haploytpes have more extreme effects for (A) eQTL and (B) cQTL compared with the other strains.** Haplotype effects were estimated as BLUPs, which are constrained and centered around 0. Each QTL is represented by an 8-element effect vector. Founders with more extreme effects are identified by comparing the absolute values of effects. Founder haploytpe effect trends for eQTL are similar to cQTL. The trends are unstable in distal-cQTL because so few are identified.

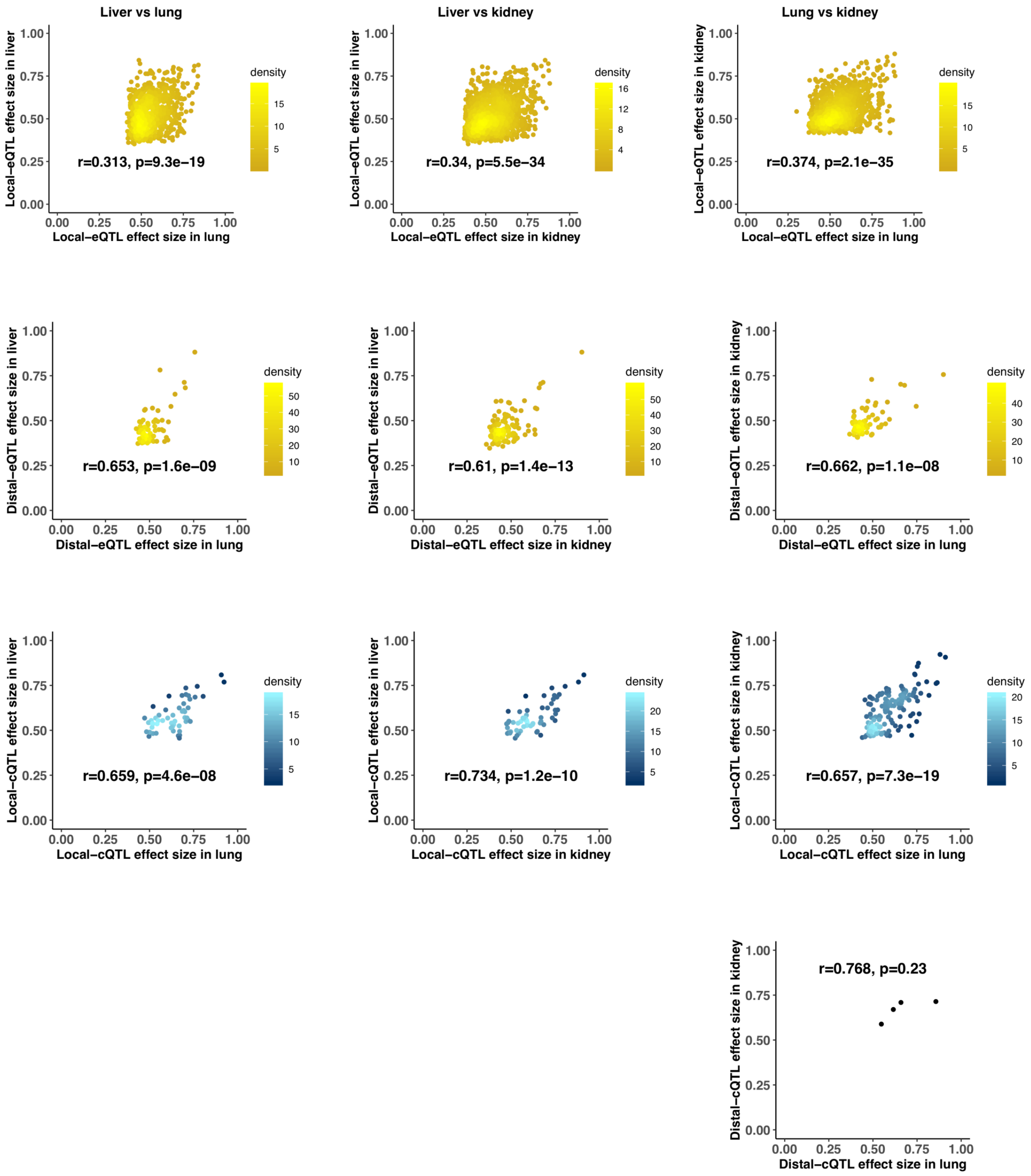


Fig S9: **Effect sizes between cross-tissue QTL pairs are lowly but significantly correlated.** Comparisons of QTL effects sizes between (liver/lung) are in the left column, (liver/kidney) middle column, and (lung/kidney) right column. eQTL are yellow and cQTL are blue. Local-eQTL are plotted in the top row, distal-eQTL in the second row, local-cQTL in the third row, and distal-cQTL in the bottom row, with only four pairs detected in (lung/kidney).

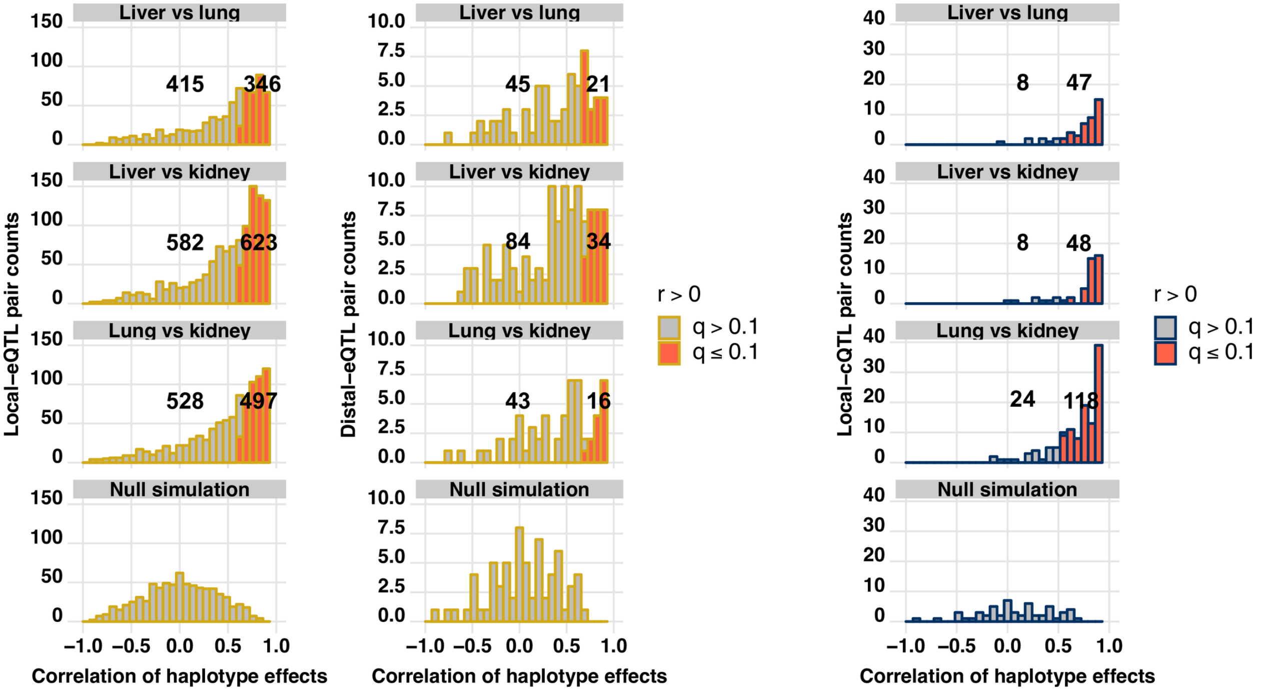


Fig S10: **Consistent genetic regulation of gene expression and chromatin accessibility observed across tissues.** There was an excess of significant positively correlated haplotype effects in QTL pairs across tissues for gene expression and chromatin accessibility. Pairs of QTL observed in multiple tissues were defined for local-eQTL (left column), distal-eQTL (middle column), and local-cQTL (right column). Only four pairs of distal-cQTL were observed, all shared between lung and kidney. A right-tailed test of the correlation between haplotype effects ($H_A : r > 0$) was performed for each QTL pair, producing $p$-values that were then FDR adjusted. Null simulations of uncorrelated 8-element vector pairs for each class of QTL and pairwise tissue comparison emphasize the observed enrichment in correlated haplotype effects between QTL pairs.

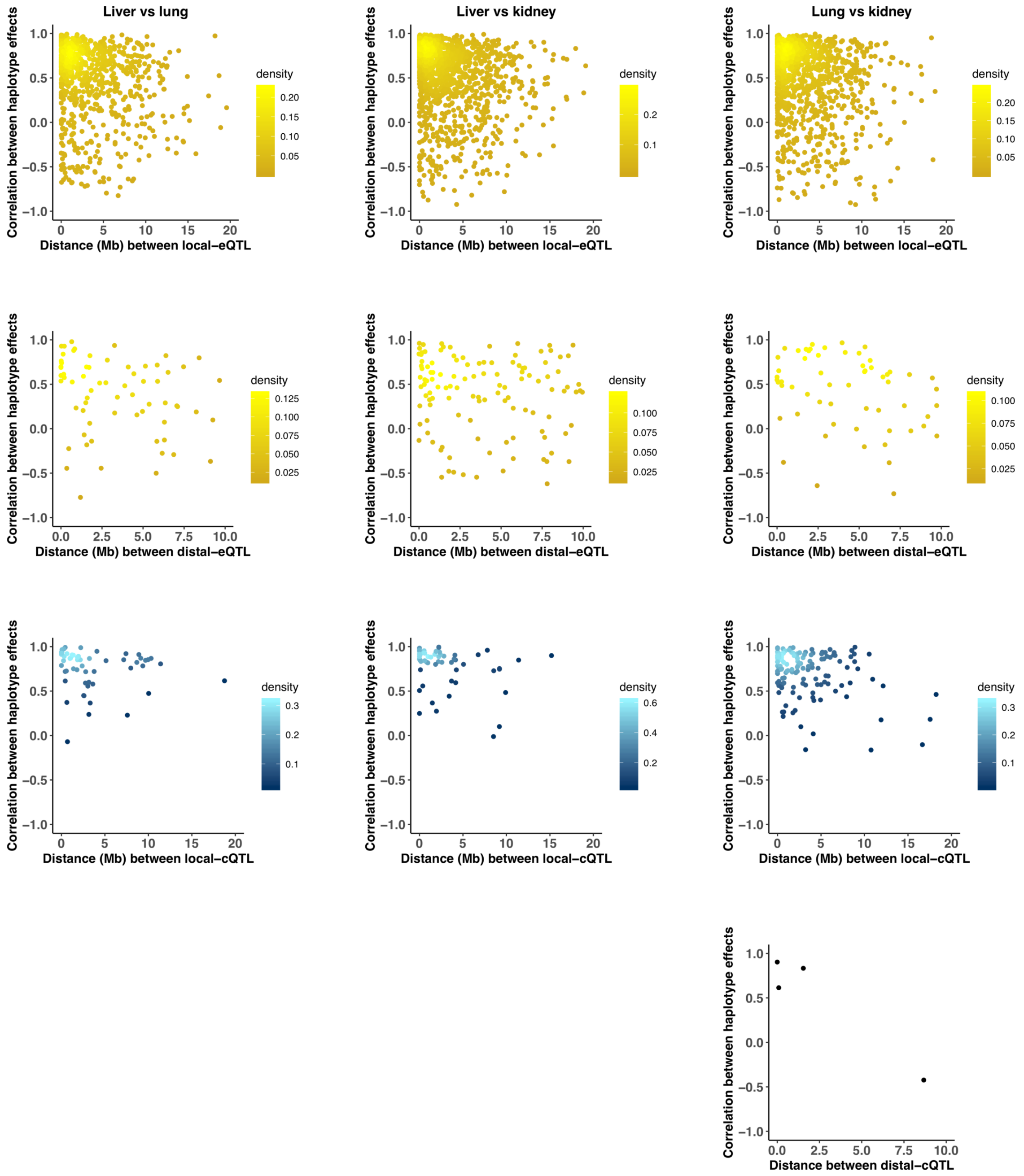


Fig S11: **Cross-tissue QTL pairs with highly correlated haplotype effects map proximally to each other.** Haplotype effects were estimated as constrained BLUPs. Pairwise correlations of the 8-element effect vectors were calculated for QTL pairs, and plotted again the distance between the QTL coordinates inMb for (liver/lung) in the left column, (liver/kidney) in the middle column, and (lung/kidney) in the right column. Single eQTL and cQTL pairs are represented as a yellow and blue dots, respectively. Local-eQTL are shown in the top row, distal-eQTL in the second row, local-cQTL in the third row, and distal c-QTL in the bottom row, for which only four pairs were detected in (lung/kidney).

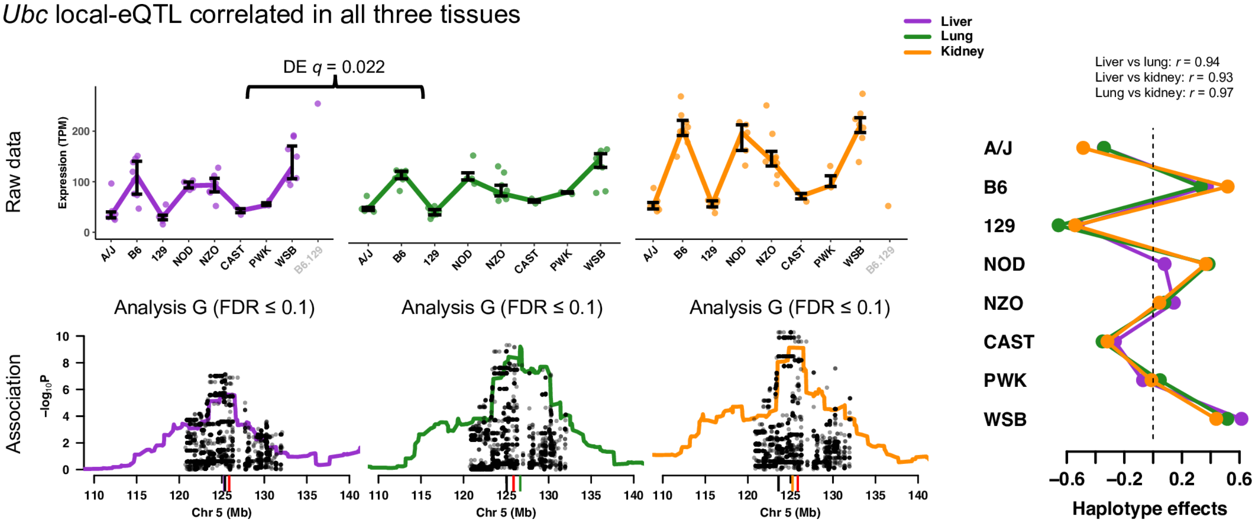


Fig S12: **The gene Ubc has consistent strong local-eQTL observed in the three tissues.** The local-eQTL consistently drove higher expression when the B6, NOD, NZO, and WSB haploytpes were present. Expression levels in liver and lung were found to be significantly different ($q = 0.022$). The estimated haplotype effects were highly consistent with the expression data, represented as interquartile bars categorized by most likely diplotype. The haplotype and variant associations in the eQTL regions were similar across tissues, suggesting they may represent the same causal origin. The red tick represents the *Ubc* TSS, the black tick represents the peak variant association, and the colored ticks represent the peak haplotype association for each tissue.

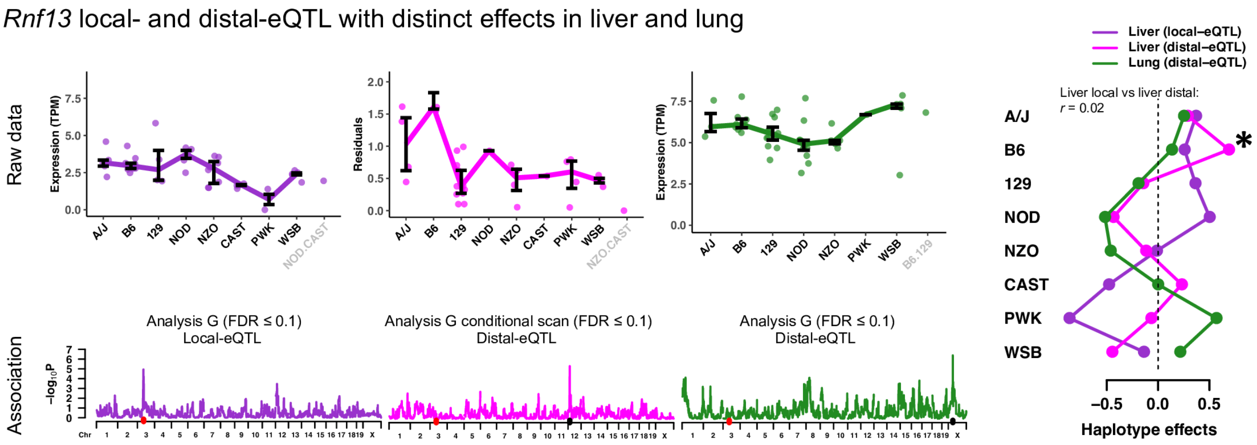


Fig S13: **The gene Rnf13 has unique patterns of genetic regulation across tissues.** A strong local-eQTL was detected in liver, and after conditioning on it, a statistically significant distal-eQTL was detected (Analysis G) on chromosome 12, largely driven by the B6 haplotype, distinct from the local-eQTL. The unique haplotype effect patterns for each eQTL can be seen in both the expression data, represented by interquartile bars for most likely diplotype, and the estimated effects. The red tick marks the *Rnf13* TSS and the black tick marks the location of distal-eQTL. Another strong distal-eQTL was detected on the X chromosome in lung.

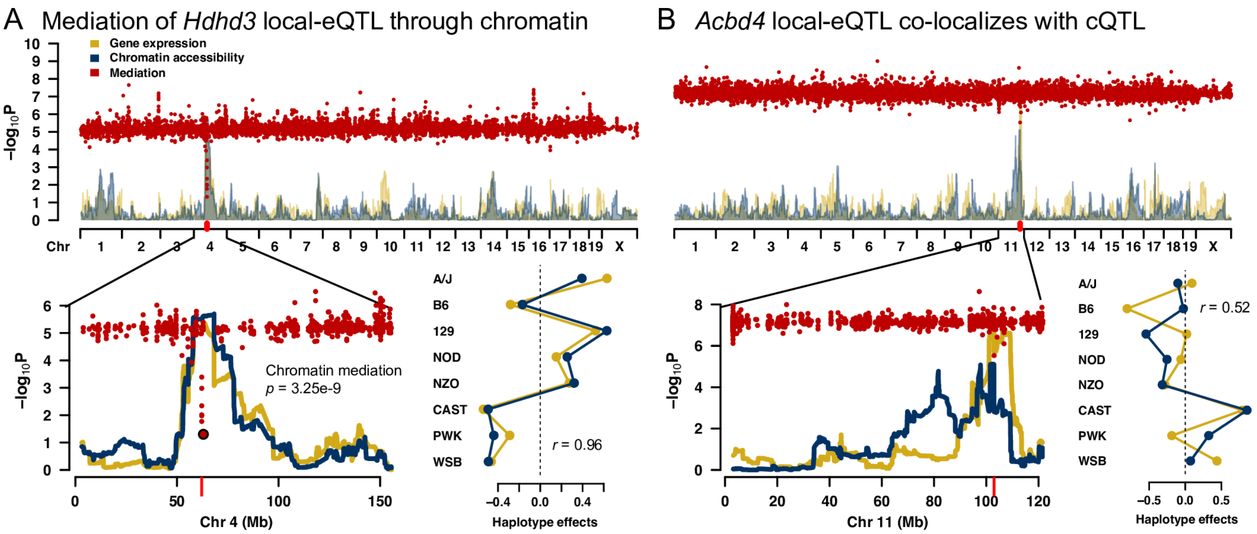


Fig S14: **Co-localizing eQTL and cQTL are not sufficient for statistical mediation.** The approach used to detect mediation through chromatin accessibility requires that the eQTL and cQTL co-localize (both within 10Mb of the gene TSS), as well as possess similar haplotype effect patterns. Co-localizing cQTL are observed for local-eQTL for both (A) *Hdhd3* in liver and (B) *Acbd4* in kidney. QTL and mediation scans are shown, with chromosomes 4 and 11 blown up for *Hdhd3* and *Acbd4*, respectively. The red ticks denote the TSS for both genes. The haplotype effects for the eQTL and cQTL are highly correlated ($r = 0.96$) for *Hdhd3*, but not for *Acbd4* ($r = 0.55$). Strong mediation of the *Hdhd3* eQTL through chromatin is detected, but not for *Acbd4*. The effect size of the co-localizing cQTL to *Acbd4* is smaller than its eQTL, also inconsistent with the relationship depicted in Fig 6A.

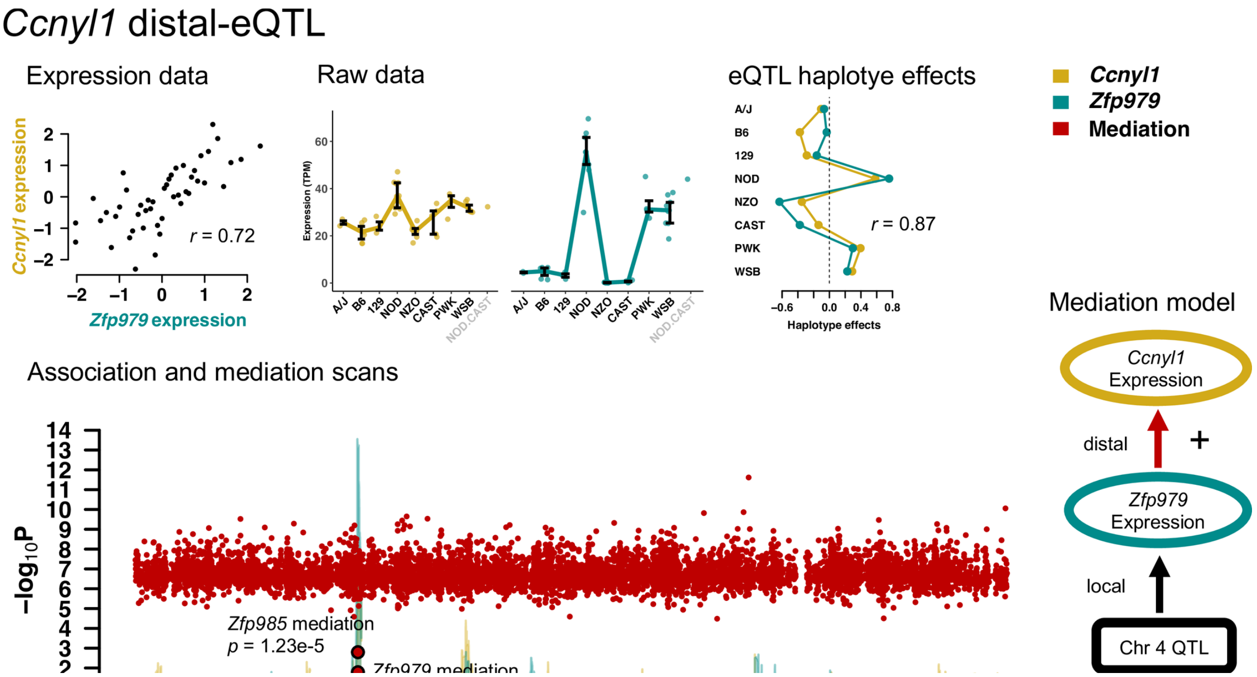


Fig S15: **Mediation of Ccnyl1 distal-eQTL through Zfp979 expression.** Expression of *Ccnyl1* and *Zfp979* are correlated $(r = 0.72)$ in lung, which is also observed in the expression data categorized by diplotype and the haplotype effects The distal-eQTL on chromosome 4 for *Ccnyl1* corresponds closely to local-eQTL of *Zfp979*. *Ccnyl1* is located on chromosome 1, indicated by the red tick. *Zfp979* and *Zfp985*, both zinc finger proteins likely with DNA binding properties, are identified as strong candidate mediators of the distal-eQTL at genome-wide significance. The correlations, magnitude of effects, and mediation are consistent with the simple relationship depicted in the graph. The distal-eQTL and candidate mediators are located in a region of interest that regulates *Akr1e1* expression.

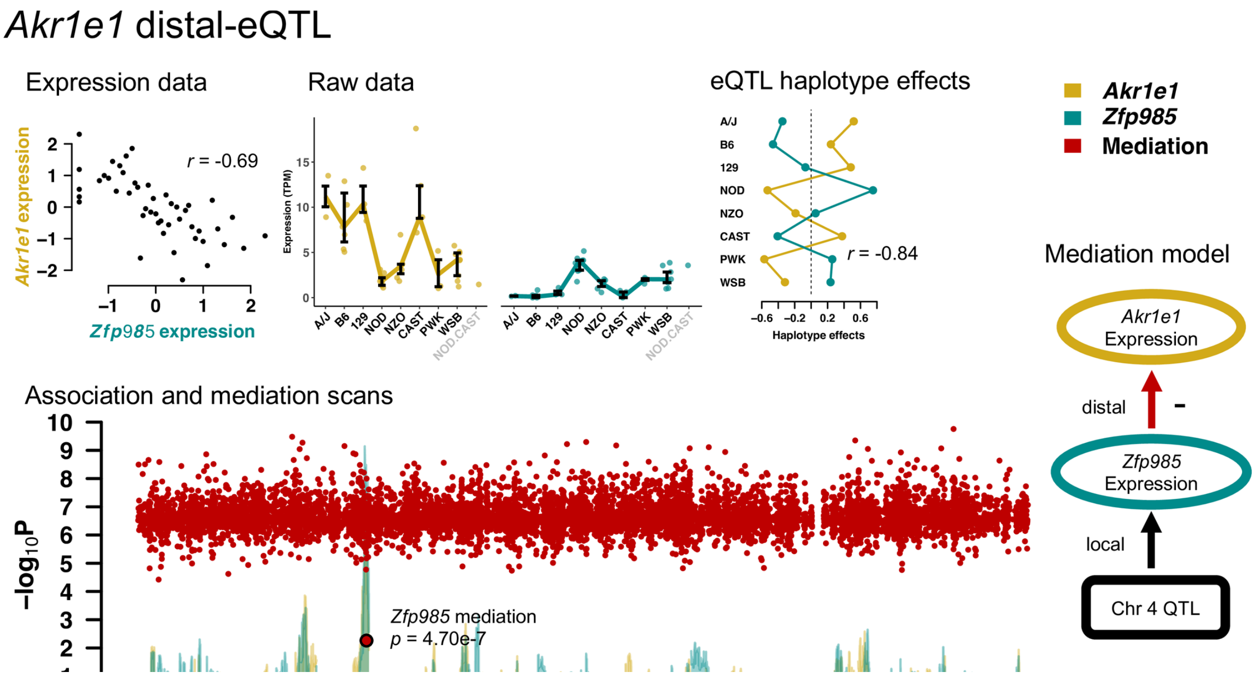


Fig S16: **Mediation of Akr1e1 distal-eQTL through Zfp985 expression.** Expression of *Akr1e1* and *Zfp985* are anti-correlated ($r = -0.69$) in lung. This relationship is also observed in the expression data with bars representing the interquartile range, categorized by most likely diplotype, and the haplotype effects. The QTL and mediation scans reveal that *Akr1e1*, with TSS marked with a red tick on chromosome 13, possesses a distal-eQTL on chromosome 4 that is nearby the strong local-eQTL of *Zfp985*. The mediation scan identifies *Zfp985* as a strong candidate mediator consistent with the mediation model. A more complete picture of the genetic regulation of *Akr1e1* expression is pieced together by looking across all three tissues and includes a potential chromatin mediator (Fig 9).

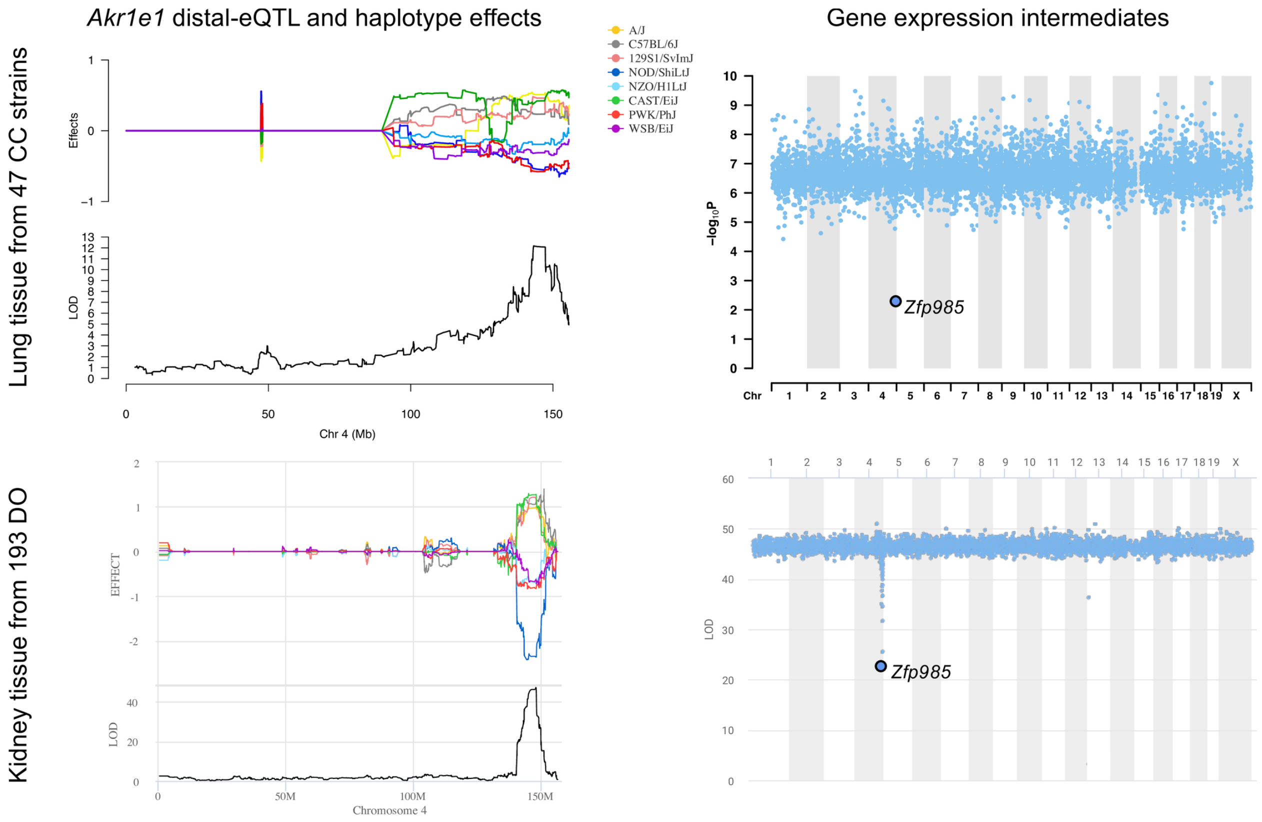


Fig S17: **Confirmation of Akr1e1 distal-eQTL and mediation by Zfp985 in kidney tissue of Diversity Outbred mice.** A genome-wide significant distal-eQTL was detected for *Akr1e1* in liver, lung (shown here), and kidney tissues from 47 CC strains. In a larger sample of kidney tissue from outbred DO mice, the same distal-eQTL and mediation relationship were observed. As expected, the larger sample of the DO results in greater statistical significance, and confirms that the NOD effect is more strongly negative than NZO, PWK, and WSB, which the haplotype effects plots for the *Zfp985* local-eQTL suggested. Notably, *Zfp985* was not tested in the CC kidney because of low expression levels, though the distal-eQTL for *Akr1e1* is consistent with its activity, which is here confirmed in the DO.

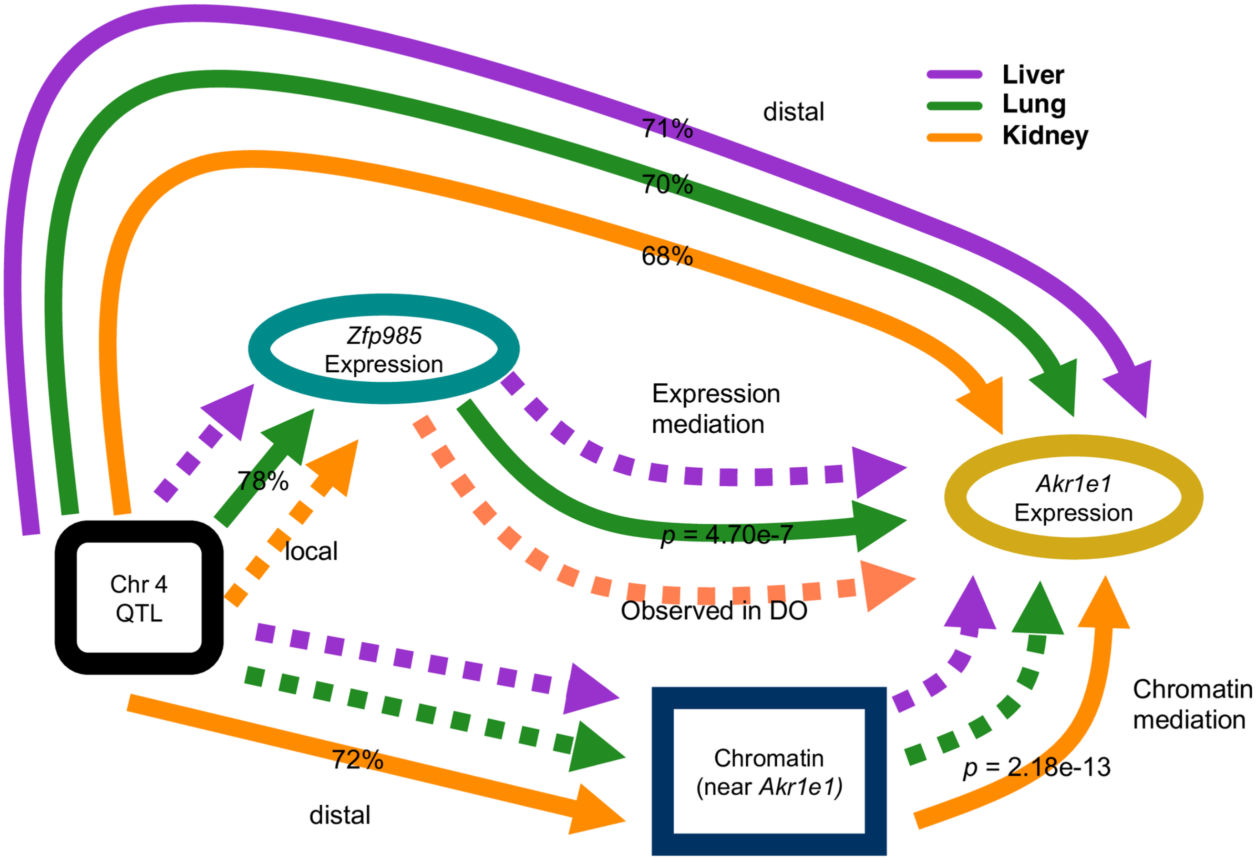


Fig S18: **Observed relationships across the three tissues related to the genetic regulation of Akr1e1 expression.** The model for the distal genetic regulation of *Akr1e1* expression, described in Fig 9, was reconstructed from these observed relationships. Solid arrows were observed, whereas dashed arrows are assumed. QTL effect sizes represent the proportion of variance explained by the QTL and mediation $p$-values (permP) were defined using a permutation procedure. The assumed relationships are supported by the presence of the distal-eQTL in all three tissues. The *Zfp985* mediator relationship in kidney, though not observed in the CC, was observed in the related DO population.

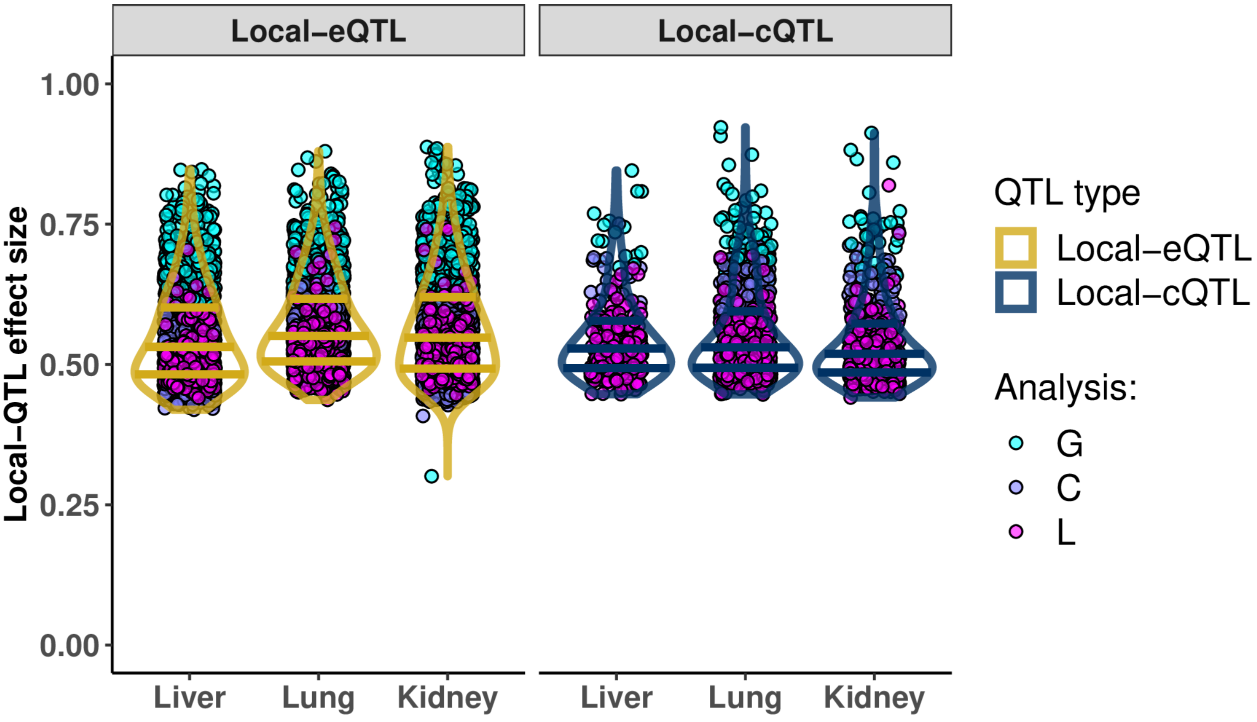


Fig S19: **Local-QTL effect sizes by mapping analysis.** Based on ranking mapping analyses with respect to the extent of scope, local (L; magenta) to chromosome (C; plum) to genome-wide (G; cyan), the greater the scope corresponded to reduced power to detect QTL, shown in liver, lung, and kidney tissues for gene expression (yellow line) and chromatin accessibility (blue line). Each dot represents a detected local-QTL, colored according to the highest scope mapping procedure that detected it. The three horizontal bars represent the $25^{th}$, $50^{th}$, and $75^{th}$ quantiles of QTL effect sizes for all local-QTL per tissue. Analysis G generally detects QTL with effect size $> 60\%$, whereas Analyses C and L detect QTL effect sizes $> 45\%$. Effect size estimates correspond to a fixed effects model of the QTL.

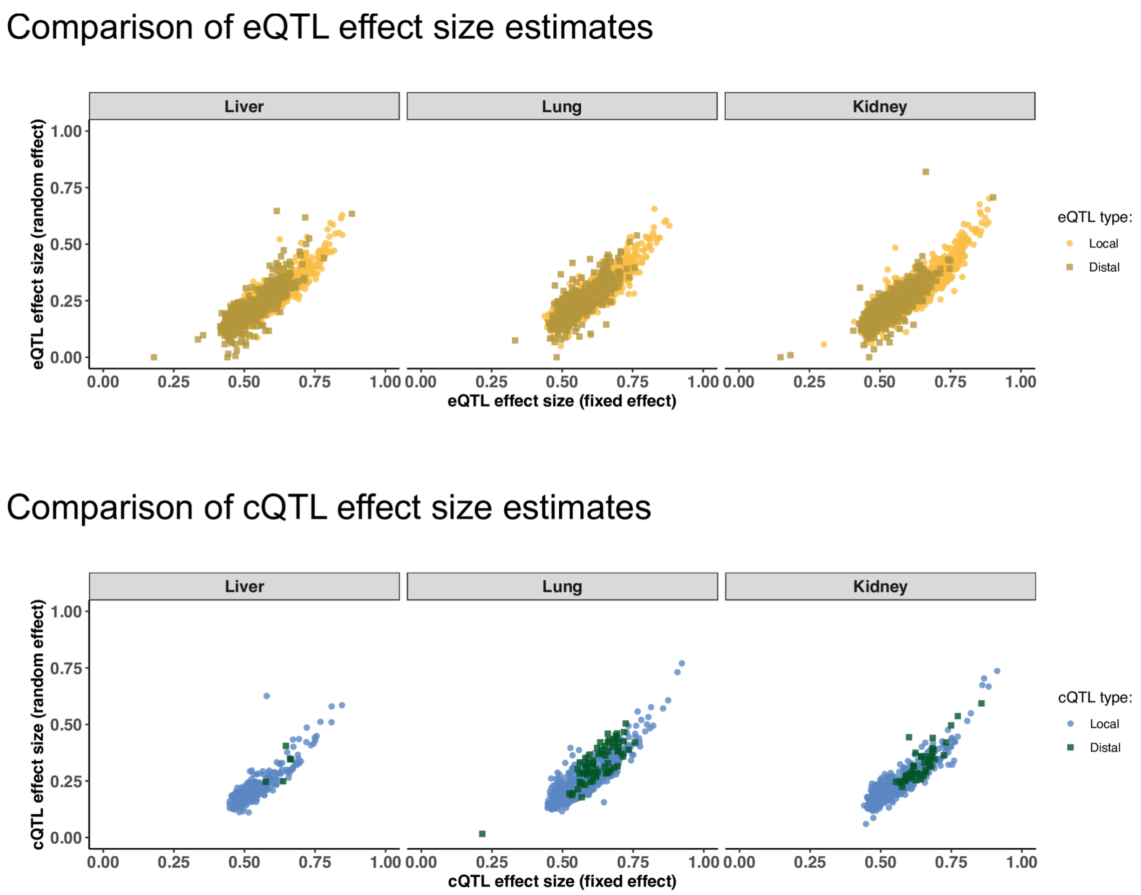


Fig S20: **Comparison of QTL effect sizes estimates from fixed effects and random effects models.** The effect size from the the random effect fit is harshly penalized compared to the fixed effect estimate, likely due to a sample size of 47 mice. Notably, there are a number of distal-eQTL that are more harshly reduced by the random effects model compared to the other QTL, likely representing signals resulting from extreme observations or imbalances in founder contributions at the locus. QTL detected by Analyses G (FDR $\leq 0.1$), C (FDR $\leq 0.1$), and L are shown.

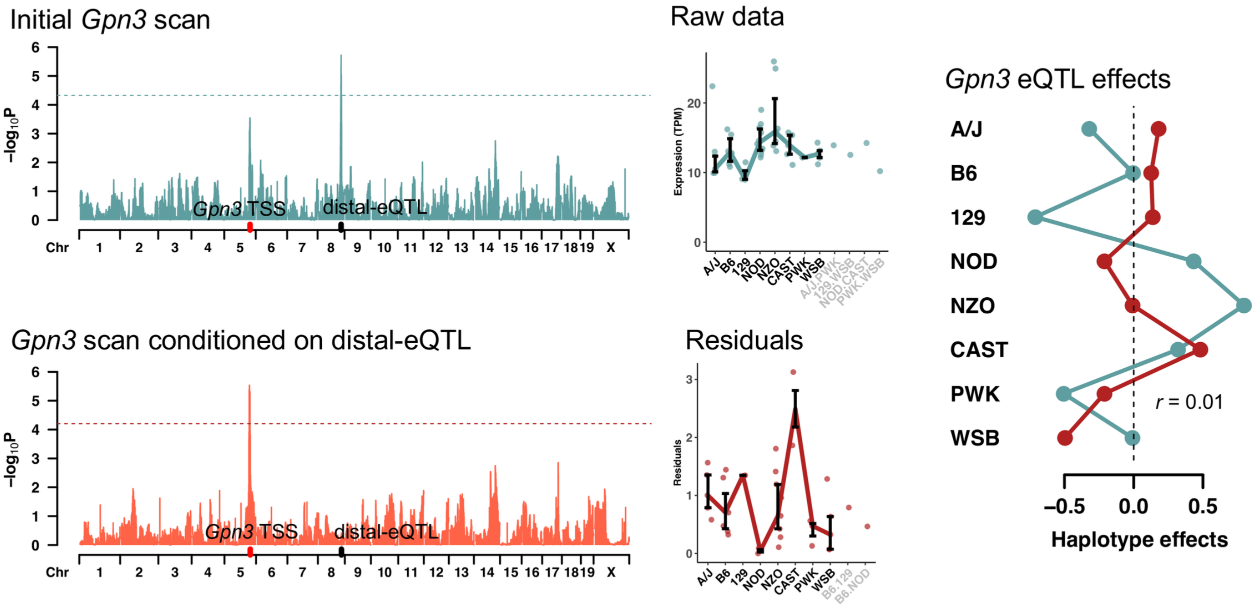


Fig S21: **Detection of local-eQTL after conditioning on distal-eQTL for Gpn3.** The multi-stage conditional regression approach of Analysis G allows for the detection of multiple genome-wide significant QTL, which can then be appropriately incorporated into an FDR procedure across many outcomes. In this example in lung tissue, the gene *Gpn3* initially has a strong distal-eQTL on chromosome 8 [top left]. Though a peak is detected near the TSS of *Gpn3*, it does not meet genome-wide significance. However, after conditioning on the distal-eQTL, the local-eQTL is detected [bottom left]. Horizontal dashed lines represent empirical 95% significance thresholds based on 1,000 permutations.
