## Supplementary material for "Integrative QTL analysis of gene expression and chromatin accessibility identifies multi-tissue patterns of genetic regulation": File S24

### S1 Appendix: CC strains

This study included a single male from the following 47 CC strains: CC001, CC002, CC003, CC004, CC005, CC006, CC007, CC010, CC011, CC012, CC013, CC015, CC016, CC017, CC019, CC020, CC021, CC023, CC024, CC025, CC027, CC028, CC029, CC030, CC031, CC032, CC033, CC035, CC036, CC037, CC038, CC039, CC040, CC041, CC042, CC043, CC044, CC045, CC046, CC049, CC053, CC055, CC057, CC059, CC061, CC062, and CC068.

### S2 Appendix: Detailed description of conditional genome-wide scans (Analysis G)

QTL analysis for a large panel of molecular traits, such as gene expression or chromatin accessibility, poses a large multiple testing problem: all loci tested for each of many traits. Analysis G employs a per-trait family-wise error rate (FWER) control and across-trait false discovery rate (FDR) control, which restricts its ability to detect multiple QTL per outcome in a non-biased manner. This bias is due to the generalized extreme value distribution (GEV), used for FWER control, being fit from the maximum statistical score per trait. To include additional strong statistical scores is potentially biasing, as would be the case when there are multiple associations above the FWER threshold, increasing the enrichment in small $p$-values, which is precisely the pattern of association FDR procedures aim to detect. To avoid this problem, a multi-stage conditional regression approach was used [1]. The procedure is described in the following steps:

1. For a given trait, conduct a genome scan according to model:

$$y_i = \mu + \text{batch}_{b[i]} + \text{QTL}_i + \varepsilon_i, \tag{1}$$

   where $y_i$ is the trait level for individual $i$, $\mu$ is the intercept, $\text{batch}_b$ is a categorical fixed effect covariate with five levels $b = 1, \ldots, 5$ representing five sequencing batches for both gene expression and chromatin accessibility and $b[i]$ denoting the batch relevant to $i$, $\varepsilon_i \sim \text{N}(0, \sigma^2)$ models the residual error, and $\text{QTL}_i$ models the genetic effect at the locus.

2. Perform permutation scans to characterize a GEV. Calculate a genome-wide error rate controlled $p$-value, $\text{permP}_\text{G}$, from the observed maximum logP of the genome scan. The $\text{permP}_\text{G}$ is stored to be used as input to an FDR procedure.

3. Specify a genome-wide $\alpha_\text{step}$ for determining whether subsequent conditional scans should be conducted for the trait. We set $\alpha_\text{step} = 0.1$. If $\text{permP}_\text{G} > \alpha_\text{step}$, no further conditional scans are conducted.

4. If $\text{permP}_\text{G} \leq \alpha_\text{step}$, steps 1-3 are repeated for an additional conditional scan of the outcome. For $j > 1$, the $j^\text{th}$ conditional scan uses the same form of alternative and null model as described in Eq 1, except with the inclusion of locus effects from the peak associations from previous stages. Generally, the alternative model for conditional scan $J$ follows as:

$$y_i = \mu + \text{QTL}_i + \sum_{j=1}^{J-1} \text{locus}_i^j + \text{batch}_{b[i]} + \varepsilon_i, \tag{2}$$

   with $\text{locus}_i^j$ representing the locus effect of the peak association from the $j^\text{th}$ stage scan of the outcome for individual $i$, and is also included in the null model of conditional scans. Now steps 2-4 are repeated.

A multi-stage conditional scan approach could have issues with over-fitting, given the data comprise 47 mice and each QTL and locus effect represents the estimation of seven fixed effects. However, permutation thresholds were found to appropriately compensate for this potential problem based on the recalculation of the GEV using permutations of the conditional scans in step 2. Fig S21 provides a clear example in which the gene *Gpn3* has a local-eQTL detected after a strong distal-eQTL is conditioned on in lung tissue.

### S3 Appendix: Mediation analysis

For this study, simple mediation models consisting of three variables were used: the independent variable X, the mediator M, and the dependent variable Y [2, 3]. Superscripts are used to indicate whether a variable is the QTL, chromatin accessibility at site $k$ ($\mathrm{C}_k$), or expression of gene $j$ ($\mathrm{G}_j$) in the model. When an eQTL is detected, the following relationship is supported:

$$\mathrm{X}^{\mathrm{QTL}} \quad \rightarrow \quad \mathrm{Y}^{\mathrm{G}_j} \tag{3}$$

Candidate mediators of eQTL, either nearby chromatin regions or genes, must also possess their own local-QTL, represented similarly:

$$\mathrm{X}^{\mathrm{QTL}} \quad \rightarrow \quad \mathrm{M} \tag{4}$$

where M is some $Y^{G_k}$ or $Y^{C_k}$, a candidate mediator of the eQTL of $Y^{G_j}$. Here we consider two models of mediation: 1) the effect of an eQTL is mediated through local chromatin state ($\mathrm{M}^{C_k}$) and 2) the effect of the distal-eQTL is mediated through a nearby gene ($\mathrm{M}^{G_k}$), depicted in Fig 6. Co-localizing QTL are consistent with the co-occurrence of $X \rightarrow Y^{G_j}$ and $X \rightarrow Y^{C_k}$ at a locus. If X is fully independent of Y given M, denoted as:

$$\mathrm{X}^{\mathrm{QTL}} \quad \perp\!\!\!\perp \quad \mathrm{Y}^{\mathrm{G}_j} \quad | \quad \mathrm{M} \tag{5}$$

where "A $\perp\!\!\!\perp$ B" indicates that A and B are independent of each other and "A | B" is A conditioned on B, then X acts on Y completely through M, which is commonly referred to as full mediation. Partial mediatory relationships are also possible in which X affects Y both directly and through M, traditionally identified by testing for:

$$\mathrm{M} \quad \rightarrow \quad \mathrm{Y}^{\mathrm{G}_j} \quad | \quad \mathrm{X}^{\mathrm{QTL}} \tag{6}$$

In practice, the true relationships among the variables can be complex and obscured by noise, making it hard to completely distinguish full mediation, partial mediation, and X separately affecting M and Y. Furthermore, genome-wide data present a significant challenge to systematically testing these relationships because of the presence of many candidate mediators, potential correlation structure due to linkage disequilibrium (LD), and error heterogeneity across the levels of genomic data. For these reasons, we do not use the traditional mediation tests [2], but adapt and simplify them for genomic data.

#### Scans

We perform mediation scans, described in Eq 3 of the Methods. These scans test for significant reduction in the relationship $X \rightarrow Y|M$, akin to a full mediator (relationship 5), though without attempting to define a formal threshold for defining whether variables are weakly associated or independent. The relationship is tested for all mediators within the local region of the eQTL (±10Mb), thus could plausibly explain the eQTL.

##### Permutations

Empirical $p$-values for the mediators can be defined through permutations, similarly to as is done with the QTL. Instead of permuting the trait outcome, the pairing of trait to QTL is maintained, and the mediator is permuted. GEVs are then fit for potential mediators from the minimum logP from the permutation scans, instead of the maximum logP in QTL scans. FWER-controlled mediation $p$-values are calculated from the cumulative distribution function of the GEV: $\mathrm{permP}^m = F_{\mathrm{GEV}}(\mathrm{logP})$. An additional FDR adjustment is not used with mediation because it is only evaluated for detected QTL, which we found likely to violate the uniform null distribution expectations of FDR. Notably, we use the genome-wide and chromosome-wide measurement of molecular traits, the vast majority of which are ineligible to be mediators of the eQTL due to location, to characterize empirical error distributions for the mediator test, and thus derive FWER-controlling thresholds. This approach is statistically stringent, given that mediators must be local to the eQTL, which is appealing in terms of controlling false positives that could result in biologically-complex data.

#### Mediation criterion

##### Mediation of local-eQTL through chromatin state.

Mediation through chromatin accessibility was evaluated for all local-eQTL detected through the lenient Analysis L, leveraging prior knowledge of the predominance of local genetic regulation and the effects of proximal chromatin state. Candidate mediators are detected at genome and chromosome-wide significance through 1,000 permutations. Genome-wide mediation is detected if $\text{permP}_{\text{G}}^{m} < 0.05$ and chromosome-wide if $\text{permP}_{\text{C}}^{m} < 0.05$.

##### Mediation of distal-eQTL through local gene expression.

Mediation of the effect of genome-wide significant distal-eQTL (Analysis G) through the expression of nearby genes was evaluated. Genome-wide significance is required for distal-eQTL because there is not additional strong biological information to support reduced stringency as is the case with local-eQTL. As with mediation through chromatin state, 1,000 permutations are used to derive genome-wide mediation $p$-values, $\text{permP}_{\text{G}}^{m}$.

Mediation of cQTL through other chromatin sites and gene expression were also considered, but did not result in the detection of any candidate mediators other than *Zfp985* expression as a mediator of a distal-cQTL on chromosome 4 for a chromatin region located chromosome 13 near *Akr1e1*.

##### Mediation steps

There are important caveats and pitfalls for mediation analysis compared to QTL mapping, in part due to the fact that the relationships 3 and 4 could be complex, whereas the directionality of the relationships for QTL is fixed because the genetic state at the QTL cannot change because of the trait. This does not hold for the mediation model, where Y could in fact causally influence M ($\text{Y} \rightarrow \text{M}$). False mediation can be detected when the association between X and M are particularly strong, whereby M acts as a surrogate for X [4]. Based on concerns over fitting relatively complex models to data comprising 47 mice, we use stringent requirements for declaring evidence for mediation:

1. Detection of a significant eQTL (relationship 3).
   (a) Mediation through chromatin: $\text{permP}_{\text{C}}^{m} < 0.05$ from Analysis L
   (b) Mediation through expression: $\text{permP}_{\text{G}}^{m} < 0.05$ from Analysis G
2. Detection of significant mediator within a $\pm$10Mb window around the eQTL from previous step.
3. Detection of a significant QTL for the mediator from previous step.
   (a) Mediation through chromatin: $\text{permP}_{\text{C}}^{m} < 0.05$ from Analysis L and within $\pm$10Mb of gene TSS
   (b) Mediation through expression: $\text{permP}_{\text{G}}^{m} < 0.05$ from Analysis G and within $\pm$10Mb of the eQTL position
4. It is not possible to distinguish $\text{M} \rightarrow \text{Y}$ from $\text{M} \leftarrow \text{Y}$ given these data. If M is an intermediary of X on Y, the expectation is that the effect of $\text{X} \rightarrow \text{M}$ is greater than $\text{X} \rightarrow \text{Y}$. We check that the effect size of the QTL for step 1 ($\text{X} \rightarrow \text{Y}$) is less than the effect size of the QTL from step 3 ($\text{X} \rightarrow \text{M}$). Failure of this step does not disprove the proposed mediation model, but it could suggest that the relationship between M and Y is more complicated than the simple models tested here, or that greater noise is incurred in the measurement of M than Y.

Note: Step 4 is an approximation of testing whether M provides more information about Y than X, which in theory, could be more directly tested with the partial mediation test (relationship 6). However, we found this test was inconclusive (partial mediation $p$-value $< 0.1$) for the clear example of *Zfp985* expression mediating the distal-eQTL of *Akr1e1*, suggesting the error heterogeneity across the measurements of these molecular phenotypes may be too great to allow for full use of traditional mediation tests, such as distinguishing partial mediation, full mediation, and X affecting M and Y separately.
